# Cultivar- and field site-specific protein and metabolite patterns of faba bean (*Vicia faba* L.) seeds

**DOI:** 10.1101/2025.10.14.682483

**Authors:** Julia Brandt, Alexandra Hüsken, Eileen Whitelaw, Georg Langenkämper, Rabea Schweiger

**Affiliations:** Department of Chemical Ecology, Bielefeld University, Bielefeld, Germany; Max Rubner-Institut (MRI) - Federal Research Institute of Nutrition and Food, Department of Safety and Quality of Cereals, Detmold, Germany

**Keywords:** cultivar, faba bean, field site, metabolites, metabolic fingerprinting, proteins, seed quality

## Abstract

Sustainable agriculture is an important component of current and future food production, particularly in the face of the rising human world population. In this context, faba bean (*Vicia faba* L.) is highly valuable due to its capability to enrich soils with nitrogen and because its seeds have a high nutritional value. In this study, we compared the seed chemistry of 11 faba bean cultivars grown at 13 field sites in Germany. Nitrogen and sulfur contents were determined with the Dumas method and protein contents were calculated based on the obtained nitrogen data. SDS-PAGE was used to analyze proteins, while metabolic fingerprinting was applied to screen (semi-)polar metabolites, mainly comprising specialized metabolites. The protein content differed predominantly between cultivars, but also between field sites. The sulfur content strongly depended on the field site. The patterns of proteins and metabolites were largely cultivar-specific. Additionally, the number of metabolic features varied somewhat between field sites. Our study suggests that breeding led to comprehensive changes in the seed chemistry of *V. faba* bean, going far beyond protein content and certain metabolites targeted in breeding. The rather high consistency when comparing field sites indicates that breeding has led to more or less stable cultivars. Future studies should place more emphasis on far-reaching effects of breeding on the seed proteome and metabolome of faba bean, as various proteins and metabolites in concert determine the nutritional quality of seeds of this important crop species.

## 1 Introduction

In view of the steadily increasing human world population, sustainable agriculture is crucial to fulfil the nutritional needs of the current and of future generations (Willett et al. 2019). A substantial requirement for sustainable crop production is the selection of suitable crop species and cultivars, ideally performing well under various environmental conditions at different field sites as well as under challenges associated with global change. Next to the yield of crops, their nutritional quality is of utmost importance for the consumers, being determined by compounds with beneficial as well as ones with adverse effects. Plant species within the taxon Fabaceae (“legumes”) are important in sustainable agriculture, for example in crop rotation systems, as they enrich the soils with nitrogen derived from rhizobia-mediated N_2_-fixation (Graham and Vance 2003; Jensen, Peoples, and Hauggaard-Nielsen 2010; Praharaj and Maitra 2020). In comparison to other crops, many grain legumes stand out due to high protein contents of their seeds, delivering sustainable plant-based proteins that may be used as alternatives to meat-based protein sources (Multari, Stewart, and Russell 2015; Semba et al. 2021). In addition to proteins, several primary and specialized (= secondary) metabolites are decisive for the quality of crops. Thus, it was recently highlighted that comprehensive analyses of both proteins and metabolites are needed to investigate the quality of foods, applying foodomics approaches (Balkir, Kemahlioglu, and Yucel 2021). To assess the seed quality of Fabaceae for the human diet, more studies focusing on various specialized metabolites are required (Fayek et al. 2021).

Within the Fabaceae, faba bean (*Vicia faba* L.; also called fava bean, broad bean) is an environmentally sustainable crop species used for livestock feeding and human nutrition. It enriches soils with nitrogen, promotes beneficial insects like pollinators and controls soil-borne diseases, thus being an invaluable part of sustainable cropping systems (Etemadi et al. 2019; Jensen et al. 2010).

The seeds of faba bean show a high protein content, even compared to other grain legumes, and contain various metabolites with manifold implications for the nutrition and health of livestock and humans (Crépon et al. 2010; Dhull et al. 2021; Martineau-Côté et al. 2022; Rahate, Madhumita, and Prabhakar 2021; Sharan et al. 2021; Walter et al. 2022). The daily need of essential amino acids by humans can be covered by a mixture of *V. faba* seeds with cereal grains, as both are complementary regarding their crop-specific amino acid compositions. Some of the compounds within faba bean seeds are considered to be beneficial and health-promoting, for example based on their antioxidative functions. However, there are also compounds with adverse or antinutritional effects. For example, the alkaloid (pyrimidine) glycosides vicine and convicine induce favism in humans with glucose-6- phosphate-dehydrogenase deficiency (Crépon et al. 2010). Moreover, some compounds such as phytic acid and tannins are considered to be antinutritional factors, as they lower the availability of minerals and the digestibility of nutrients. Taken together, the protein and metabolite composition of faba been seeds is decisive for their usability in livestock feeding and for human nutrition and needs to be considered in breeding and farming.

Several cultivars of faba bean are available on the market, with variation in a couple of traits, not only related to their growth and yield, but also to seed chemistry. Breeding and selection have led to cultivars differing in several variables, including yield potential and stability, resistance against certain abiotic and biotic stresses, plant growth habit, seed sizes as well as seed quality (Crépon et al. 2010; Khazaei et al. 2021; Maalouf et al. 2019; Mohammadi et al. 2024). Regarding seed chemistry, next to the desired high protein levels, efforts were undertaken to establish cultivars with reduced vicine/ convicine as well as lower tannin contents due to the adverse effects ascribed to these compounds. While some *V. faba* cultivars are mainly used for feeding of (certain) livestock, others are additionally explicitly recommended for human nutrition by the breeders. Remarkably, faba bean is cultivated in many countries around the globe and can grow under conditions that are challenging for other species including other Fabaceae species, for example under moderate waterlogging, in cold regions with a short growing period as well as in coastal regions (Etemadi et al. 2019; Jensen et al. 2010). While for certain field sites those cultivars that are recommended for such sites may be chosen, cultivars with a high yield stability in different environments may be cultivated under various abiotic and biotic conditions (Mohammadi et al. 2024).

The seed quality of *V. faba* has been assessed in several studies, often including comparisons between cultivars, growing years and/ or environmental conditions (Johnson et al. 2020; Makkar et al. 1997; Mayer Labba, Frøkiær, and Sandberg 2021; Walter et al. 2022). While the focus of many studies was on the protein content, minerals, the antioxidative capacity and/ or on certain target compounds or compound classes (e.g., vicine/ convicine, phytic acids and/ or tannins), some studies also included the protein composition and comprehensive assessments of primary and specialized metabolites. For example, Warsame et al. (2020) assessed the seed protein composition of 35 faba bean genotypes, while Shi et al. (2022) applied metabolomics to compare two genotypes. Nontargeted metabolomics studies revealed various metabolites in faba bean seeds, including primary metabolites of different classes, alkaloids, saponins, terpenoids, jasmonates and phenolic compounds (Abu-Reidah et al. 2014; Mekky et al. 2020; Valente et al. 2019). There are complex links between growth, resistance against different (a)biotic stresses, proteins and metabolism. Thus, breeding for certain desired traits, e.g., high protein content or low contents of metabolites with adverse effects, may affect the chemical composition of *V. faba* seeds far beyond these breeding targets. Likewise, various environmental factors that differ between field sites, including climatic and edaphic factors, may affect seed proteins and metabolites. More in-depth studies on the composition of proteins and metabolites in faba bean seeds including various cultivars and environmental conditions are needed to assess and optimize the nutritional quality of this promising crop species for future use in sustainable agriculture.

The aim of our study was to comprehensively compare chemical traits of *V. faba* seeds between 11 cultivars grown in one harvest year at 13 field sites distributed across Germany. Protein contents, sulfur contents, protein compositions and metabolic fingerprints, mainly comprising specialized metabolites, were evaluated to assess seed chemistry. It was expected that the seed chemistry mainly differs between cultivars mirroring breeding of distinct cultivars for specific purposes with far-reaching side effects. Moreover, based on assumed influences of varying environmental conditions, we expected some variation related to field sites.

## 2 Materials and Methods

### 2.1 Seed Material

In this study, 11 commercial spring cultivars of faba bean from two plant breeding companies (Norddeutsche Pflanzenzucht Hans-Georg Lembke, Holtsee, Germany; P. H. Petersen Saatzucht Lundsgaard, Grundhof, Germany) were included. According to the homepages of the breeders (19.01.2023), three of the varieties have low vicine/ convicine concentrations, one has a low tannin concentration, seven are mainly for livestock feeding, while the other four are additionally recommended for human nutrition. The plants were grown in 2021 on 13 agricultural field sites distributed across Germany (Figure 1), with four randomly distributed subplots (ca. 10 m²) for each cultivar per field site. These field sites are long-term study sites maintained by the Federal Office for Plant Varieties (Bundessortenamt, Hannover, Germany), applying best local agronomic practice in fertilizer and other agrochemical treatment. In this study, the cultivars are randomly numbered (C01 to C11), as the actual identities of the cultivars were of minor importance for our study and seed material was provided under the premise that cultivar names are not published. Seeds were harvested in late summer/ autumn 2021 and pooled over the subplots for each cultivar within the field sites.

**FIGURE 1.**
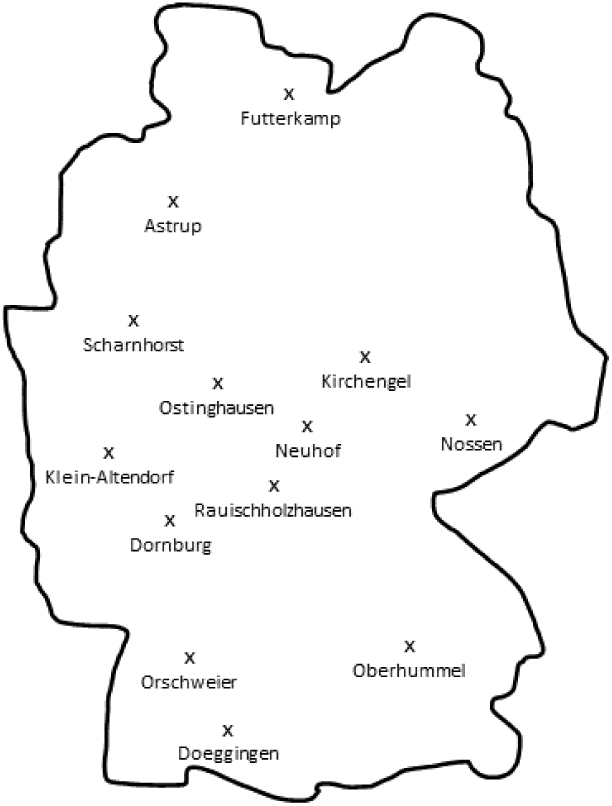
Map with the field sites maintained by the Federal Office for Plant Varieties (Bundessortenamt), from which the faba bean seeds were taken. Positions of the 13 field sites across Germany are indicated.

At the Bundessortenamt, the seeds were stored at room temperature. In May 2022, subsamples (300-400 g) were sent to the Max Rubner-Institut (Detmold, Germany), where they were stored for 2 weeks at -18 °C to kill potential pest organisms. Seeds that were very small, broken, showed mold or an unusual coloration, were shrivelled and/ or infested with *Bruchus* spp. (Coleoptera: Chrysomelidae) were discarded. Seeds were then stored at 6-8 °C until further use. Subsequent processing and analyses steps, except for protein and sulfur determination, were performed on samples that had been assembled in a randomized order. Seeds were milled in an ultra centrifugal mill (ZM 200, Retsch, Haan, Germany) using a sieve with 500 µm pore size at 14000 rpm under cooling with liquid nitrogen. Subsamples of the flour were stored at 8 °C for determination of the protein and sulfur content and at -80 °C for assessment of protein patterns and metabolite patterns, respectively. While the first three analyses were done at the Max Rubner-Institut (Detmold, Germany), the metabolic analyses were performed at the Chemical Ecology Department of Bielefeld University (Bielefeld, Germany), with an intermittent transport of subsamples on dry ice.

### 2.2 Protein Content and Sulfur Content

Nitrogen and sulfur contents of the seeds were determined via CNS elemental analysis (vario MAX cube, Elementar Analysensysteme, Langenselbold, Germany) applying the Dumas combustion principle as outlined for nitrogen (ICC-Standard 167). While the procedures of ICC-Standard 167 were followed, the combustion method using the vario MAX cube is also applicable to analyze sulfur (Schumacher, Dindorf, and Dittmar 2009). Raw protein contents were calculated by converting nitrogen contents to protein contents with the conversion factor 6.25 based on Walter et al. (2022). Protein and sulfur contents are mean values of duplicate determinations and are both given on dry mass basis.

### 2.3 Extraction and Analysis of Proteins

Seed protein patterns were analyzed in sodium dodecyl sulfate-polyacrylamide gel electrophoresis (SDS-PAGE) based on Warsame et al. (2020) with some modifications. An aliquot of 100 mg seed flour was extracted at room temperature with 1 ml of 0.1 mol L^-1^ phosphate buffer (pH 7.2) (Merck, Darmstadt, Germany) containing 28.7 mmol L^-1^ potassium sulfate (Merck, Darmstadt, Germany). Samples were vortexed (10 s) and then mixed for 1 h at 1000 rpm. After centrifugation at room temperature (17000 *g*, 30 min), supernatants were stored at -80 °C. In order to evaluate the efficiency of the extraction, a protein determination using the Dumas combustion (see 2.2) was done in triplicate on a sample extract. This analysis yielded 23.2 ± 0.1 (SD) % protein, which is 86% of the value obtained for the corresponding flour. Thus, protein concentrations in the extracts were estimated based on those in the flour, applying the correction factor 0.86. Extracts were mixed (1:1) with loading buffer [40% glycerin (86%, Carl Roth, Karlsruhe, Germany), 4% SDS (≥ 99.5%, Carl Roth), 100 mmol L^-1^ tris(hydroxymethyl)-aminomethane (TRIS; ≥ 99.9%, Carl Roth), 0.02% bromophenol blue (SERVA Electrophoresis, Heidelberg, Germany) and freshly added reducing agent (65 mmol L^-1^ 1,4-dithiothreitol, Carl Roth)], denatured (5 min, 95 °C), cooled to circa 25 °C and centrifuged (17000 *g*, 5 min). Samples (ca. 32 µg protein each) were loaded on gels [one gel per field site; separating gel: 10% polyacrylamide (with 1.5 M TRIS buffer, pH = 8.8); stacking gel: 4% polyacrylamide (0.5 M TRIS, pH = 6.8); using ammoniumperoxodisulfate (≥ 98%, Carl Roth) and tetramethylethylenediamine (Merck) for polymerisation and 10% SDS], gels were run in a PROTEAN II xi Cell (Bio-Rad Laboratories, Hercules, CA, USA) with running buffer [192 mM glycine (Carl Roth), 25 mM TRIS, 0,1% SDS; pH ∼ 8.3; 10 °C] for ca. 5 h at 32 mA (stacking gel) and 44 mA (separating gel). A protein standard (10 µL Precision Plus, 10-250 kDa, Bio-Rad Laboratories), a pool sample (originating from six extractions of seeds from the cultivar C09 and the field site Dornburg) and an “extraction-control-sample” (stemming from the same cultivar and site as the pool sample but extracted at the same time as the samples from the 11 cultivars of one field site) were additionally run on each gel. Gels were fixed [40% ethanol (96.4%, Berkel AHK Alkoholhandel, Lippstadt, Germany), 10% acetic acid (≥ 95.9%, Carl Roth), 50% Millipore-H_2_O] for 30 min and stained [0.025% Coomassie R 350 (GE Healthcare, Uppsala, Sweden) in 10% acetic acid; microwave-heated to approximately 50 °C] overnight on a shaker, destained (10% acetic acid) and scanned (GS- 800 Calibrated Densitometer, software Quantity One 4.6.9, Bio-Rad Laboratories). Gels were analyzed with Image Lab software (Bio-Rad Laboratories), ascertaining that the volumes of detected protein bands were not saturated. For relative quantification of protein bands, gel background was subtracted. Then, normalization was obtained by dividing the volume of the protein bands (area x intensity) by the volume of the most intense protein band (i.e., band no. 10 in Figure 2b) of the pool sample, yielding normalized volumes. Since the pool sample was run on each individual gel, normalization compensated for possible loading and staining differences across gels. An extraction variability of 4.4 % relative standard deviation across all 13 gels was calculated by comparing the volume of band no. 10 of the pool sample and the “extraction-control-samples” (Figures 2a,b).

**FIGURE 2.**
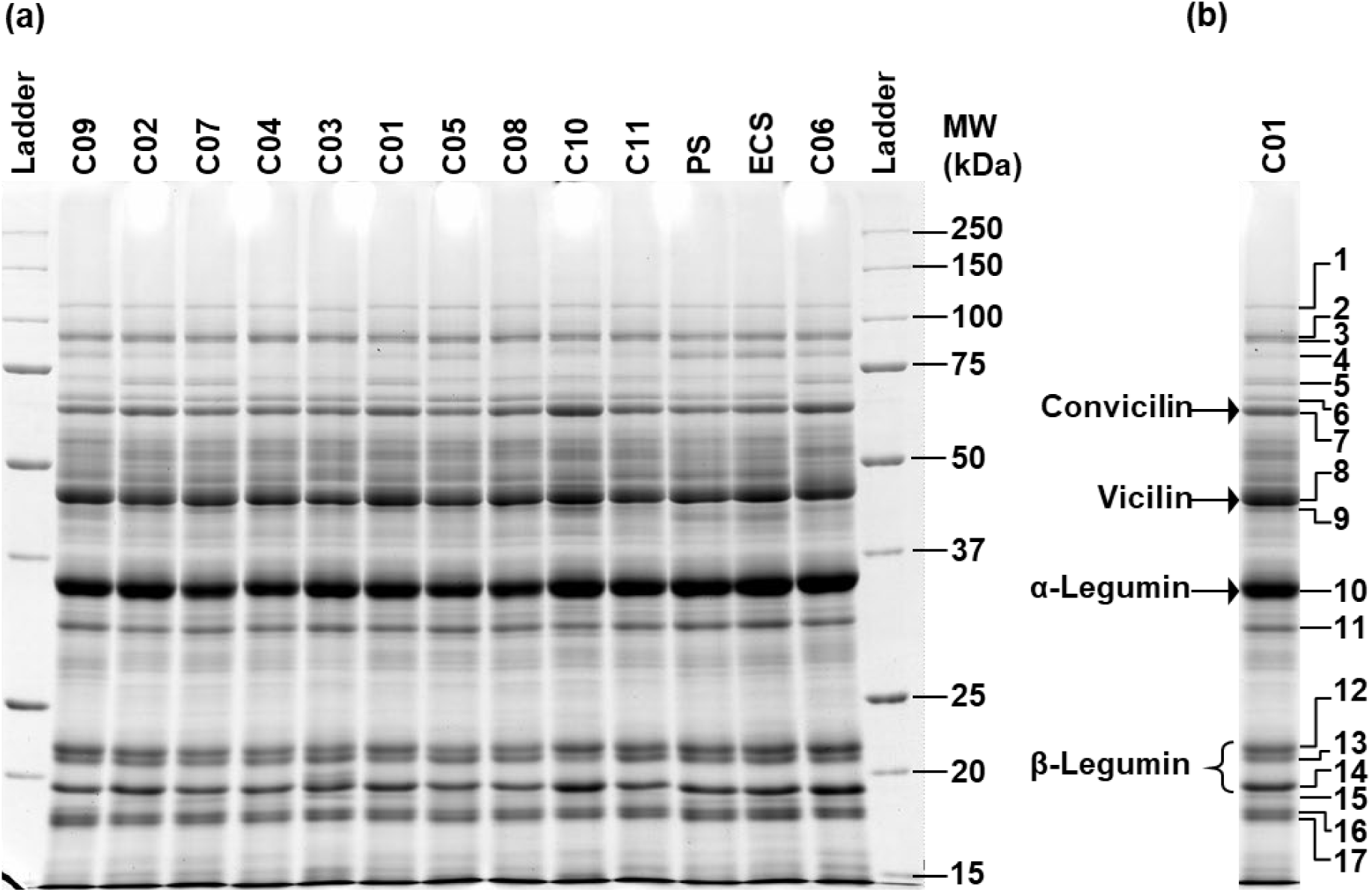
Protein pattern of 11 cultivars (C01 to C11) of faba bean seeds originating from the field site Astrup electrophoresed under denaturing conditions in SDS-PAGE (a). The pool sample (PS) and the extraction-control-sample (ECS) were used to correct for loading differences and for normalization as well as for assessing extraction variability (see Materials and Methods section). A protein standard (Precision Plus, Bio-Rad) was run to allow for estimation of molecular weights of protein bands (Ladder). (b) The single gel lane of cultivar C01 is a duplication of the same lane in (a) in order to indicate the positioning of protein band numbers (1 – 17) detected with image analysis. Based on molecular weights and comparison with literature data, the major globulin bands for convicilin, vicilin, α-legumin and β-legumin were assigned (b).

### 2.4 Extraction and Analysis of Metabolites

The metabolite patterns of the seeds were assessed by untargeted metabolic fingerprinting, mainly covering (semi-)polar specialized metabolites. Zirconium silicate beads (1.2-1.4 mm, Rimax ZS-R, Mühlmeier, Bärnau, Germany) were added (200 mg sample, 300 mg beads) on dry ice, samples stored at -80 °C and extracted on ice with 1 mL 90% methanol (LC-MS grade, Thermo Fisher Scientific, Loughborough, UK) with 0.1% formic acid (FA; RS-Pour LC-MS, CARLO ERBA Reagents, Emmendingen, Germany) and 5 mg L^-1^ hydrocortisone (Sigma-Aldrich, Steinheim, Germany) as internal standard by 10 s vortexing, homogenization (FastPrep-24 5G, MP Biomedicals, Irvine, CA, USA; 9 m s^-1^, 5 cycles of 15 s, 30 s pause time) and centrifugation (16100 *g*, 10 min). Supernatants were centrifuged again, filtered (0.2 µm, Phenomenex, Torrance, CA, USA) and stored at -80 °C. Samples were measured and data analyzed as described in Bühler and Schweiger (2024) with some modifications. Metabolites (8 µL samples) were separated by ultra-high performance liquid chromatography (Dionex UltiMate 3000, Thermo Fisher Scientific, San José, CA, USA) on a Kinetex XB-C18 column (150 x 2.1 mm, 1.7 µm, Phenomenex) at 45 °C with a flow of 0.5 mL min^-1^ and a gradient from eluent A (Millipore H_2_O with 0.1% FA) to eluent B (acetonitrile, LC-MS grade, Thermo Fisher Scientific; with 0.1% FA): 2% to 30% B within 20 min, to 75% B within 9 min, followed by column cleaning and equilibration. Part of the sample stream (split via a T-piece) was subjected to negative electrospray ionization and quadrupole time-of-flight mass spectrometry (compact, Bruker Daltonics, Bremen, Germany). Centroid data in the mass-to-charge (*m*/*z*) range of 50-1300 were recorded at 5 Hz, with the following settings (MS mode): end plate offset 500 V, capillary voltage 3000 V, low mass *m*/*z* 90, N_2_ as nebulizer (3 bar) and dry (12 L min^-1^, 275 °C) gas, quadrupole ion energy 4 eV, collision energy 7 eV. The AutoMSMS mode was used for fragmentation (MS/MS), with increasing isolation widths and collision energies along with the precursor *m*/*z*. For recalibration, sodium formate was introduced into the sprayer preceding each sample. Thirteen blanks without seed material were measured in addition. The *m*/*z* axis was recalibrated and metabolic features [each characterized by a specific retention time (RT) and *m*/*z* value] in MS mode were picked via the T-ReX 3D algorithm in MetaboScape 2021b (Bruker Daltonics). Metabolic features were picked that occurred in at least six samples with a peak height of minimum 750 and at least 11 data points. So-called buckets were created, collecting metabolic features at the same RT that probably belong to the same metabolite, using a correlation coefficient threshold of 0.8 and allowing [M–H]^−^, [M+Cl]^−^, [M+HCOOH–H]^−^, [2M–H]^−^ and [M–H–H_2_O]^−^ ions and their corresponding isotopes and charge states. From each bucket, only the metabolic feature with the highest intensity was left in the dataset and used for quantification. Metabolic features within or close to the injection peak (RT < 1.25 min) were discarded. Peak heights were divided by those of the [M+HCOOH–H]^−^ ion of hydrocortisone, background (averaged over blanks) was subtracted, features left that afterwards occurred in at least six samples and values were divided by the sample masses.

### 2.5 Statistical Analyses

All statistical analyses were done in R 4.4.2 (R Core Team 2024). Spearman rank correlations were used to test for pairwise correlations between the protein content, the sulfur content as well as the number of metabolic features. The protein content, sulfur content as well as the number of metabolic features were clustered both over cultivars and field sites (heatmap.2 function, R package *gplots*; Warnes et al. 2024). Protein and metabolite patterns were compared between cultivars and field sites using non-metric multidimensional scaling (NMDS) analyses with Wisconsin double standardization of square root-transformed data and Kulczynski distances (R package *vegan*, Oksanen et al. 2024).

Using the same data standardization and distance measure, a Mantel test with Spearman rank correlation was applied to test for correlation of protein patterns with metabolite patterns. For the protein content, sulfur content and the number of metabolic features, contour lines were plotted into the NMDS plots, using thin plate regression splines and generalized additive models (Gaussian error, identity link, restricted maximum likelihood), if the approximate smooth terms were significant (*p* < 0.05) and the explained deviance was higher than 20% (ordisurf function, R package *vegan*).

## 3 Results

### 3.1 Protein Content and Sulfur Content

The protein content of the faba bean seeds ranged from 24.0 to 33.6% dry mass (Figure 3a). It differed between cultivars and to a lower degree between field sites, being highest in the cultivar C06 as well as at the field sites Kirchengel, Rauischholzhausen and Dornburg and lowest in the cultivar C03. The seed sulfur content ranged from 0.142 to 0.233% dry mass (Figure 3b), varying mainly between field sites, with Astrup showing the highest and Scharnhorst the lowest values. The protein content and sulfur content were significantly positively correlated (Spearman correlation: *ρ* = 0.24, *p* < 0.01).

**FIGURE 3.**
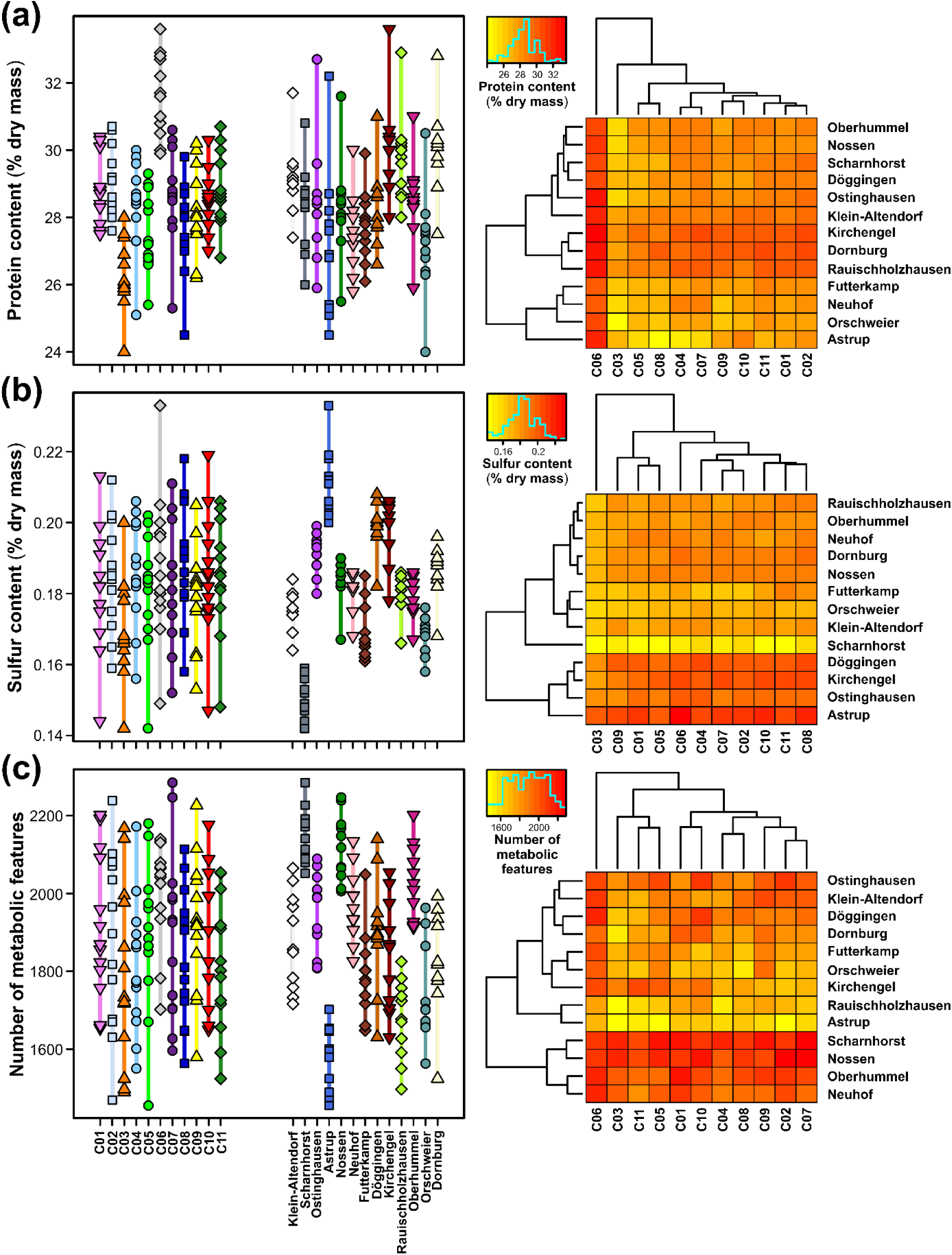
Seed chemistry of 11 cultivars of faba bean from 13 field sites. On the left, the (a) protein content, (b) sulfur content and (c) number of metabolic features are given as stripcharts, with symbol/ color coding according to cultivars (C01 to C11) and field sites, respectively. The vertical lines show the range of the data. On the right, cluster heatmaps are shown for the corresponding traits, with low values being indicated in yellow and high values in red (see color keys with histograms at top left).

### 3.2 Proteins and Metabolites

SDS-PAGE of seed proteins under reducing conditions revealed numerous protein bands, as exemplarily shown for the field site Astrup (Figure 2). The bands comprised proteins/ protein subunits with molecular weights ranging from approximately 15 kDa to 110.9 kDa. Using image analysis, in total 17 protein bands could be reliably detected, though not in all cultivars (Figure 2). Comparing the protein band pattern with published annotated SDS-PAGE images as well as an immunoblot analysis (Ashraf et al. 2020; Horstmann et al. 1993; Liu et al. 2017; Sharan et al. 2021; Tucci, Capparelli, and Costa 1991; Warsame, O’Sullivan, and Tosi 2018), the major globulin bands were assigned, i.e., convicilin (band 7) at 62.8 kDa, vicilin (band 8) at 44.3 kDa, α-legumin (band 10) at 33.8 kDa and the bands 12, 13 and 14 for β-legumin at 21.5, 20.9 and just below 20 kDa (Figure 2, Table 1). This pattern of globulin subunits matches the idealized SDS-PAGE separation of *V. faba* globulins under reducing conditions (Sharan et al. 2021). The number of detected protein bands per sample ranged from 12 to 16. Nine of these protein bands (i.e., 53%) occurred in all samples. The protein band with the lowest frequency of occurrence was band 15, which occurred in only 7 samples. The protein patterns of the seeds were compared between cultivars and field sites, using NMDS (stress value: 0.165), and strongly differed between cultivars (Figure 4a, left). For example, the cultivars C06 and C03 were distinct from each other, while several other cultivars were separated from each other as well. The cultivar C10 showed the largest variation in protein patterns. Across all cultivars, the protein bands 4, 5, 9 and 15 contributed most to the separation of the samples. In contrast to the cultivars, the protein patterns of the field sites largely overlapped (Figure 4a, right). While the NMDS ordination of protein patterns did not show obvious correlations with the sulfur content and the number of metabolic features, there was a correlation with the protein content (Figure 4a, second line). Along with the separation of the protein patterns according to the cultivars, protein contents showed a gradient from cultivar C06 with higher values to C03 and C10 with lower values.

**FIGURE 4.**
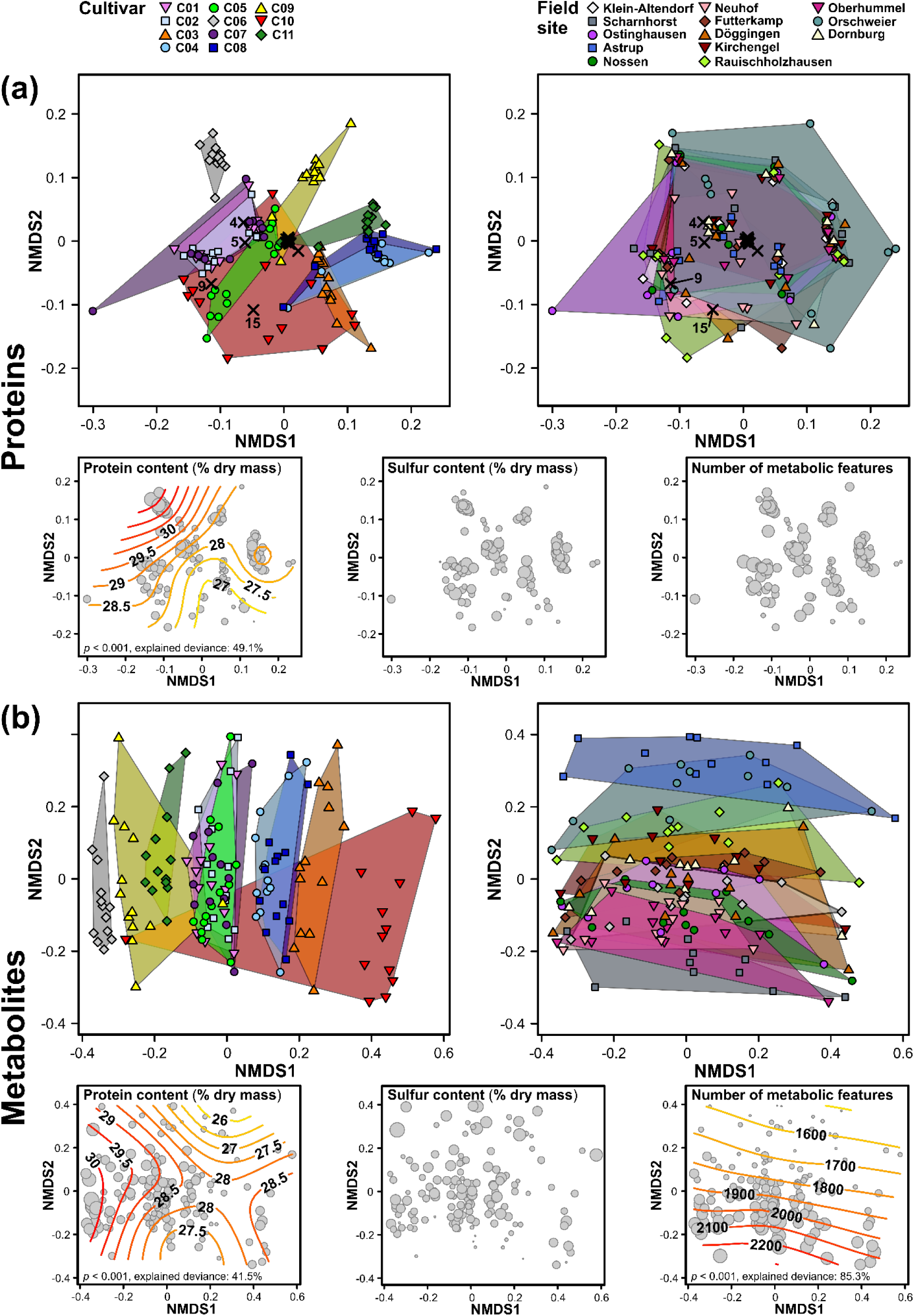
Protein and metabolite patterns of seeds of 11 cultivars of faba bean from 13 field sites. Non-metric multidimensional scaling (NMDS) analyses of (a) 17 protein bands and (b) 5598 metabolic features, with symbol/ color coding according to cultivars (left: C01 to C11) and field sites (right) and groups being surrounded by convex hulls. Loadings of proteins are indicated as crosses, with the highest loadings being labelled with the protein band number (see Table 1). The smaller plots represent simplified NMDS plots, relating the protein/ metabolite patterns to the protein content (left), sulfur content (middle) and number of metabolic features (right), with higher values being indicated by larger circles. Contour lines are shown, if the smooth terms of generalized additive models had *p* < 0.05 and an explained deviance of > 20% (given at the bottom), from low values in yellow to high values in red.

**TABLE 1.** Protein bands found per SDS-PAGE in faba bean seeds, with assignments of the major globulins, i.e., convicilin, vicilin, α-legumin and β-legumin, based on molecular weights and comparison with literature data. A typical SDS-PAGE result including positioning of protein band numbers is presented in Figure 2.

| Protein band number | Mean molecular weight (kDa) | Molecular weight (kDa) according to literature | Assignment | Literature |
| --- | --- | --- | --- | --- |
| 1 | 110.9 | - | - | - |
| 2 | 90.5 | - | - | - |
| 3 | 89.0 | - | - | - |
| 4 | 81.7 | - | - | - |
| 5 | 71.9 | - | - | - |
| 6 | 66.4 | - | - | - |
| **7** | **62.8** | **>60**<br>**~66**<br>**64**<br>**~72** | **convicilin** | Warsame et al. 2018<br>Tucci et al. 1991<br>Liu et al. 2017<br>Ashraf et al. 2020<br>Sharan et al. 2021 |
| **8** | **44.3** | **~46-55**<br>**~42-48**<br>**~55** | **vicilin** | Warsame et al. 2018<br>Tucci et al. 1991<br>Ashraf et al. 2020<br>Sharan et al. 2021 |
| 9 | 43.1 | - | - | - |
| **10** | **33.8** | **~35-40**<br>**~34-38**<br>**~35** | **α-legumin** | Warsame et al. 2018<br>Tucci et al. 1991<br>Ashraf et al. 2020<br>Sharan et al. 2021 |
| 11 | 30.5 | - | - | - |
| **12** | **21.5** | **~24**<br>**~20-25**<br>**~21** | **β-legumin** | Warsame et al. 2018<br>Tucci et al. 1991<br>Ashraf et al. 2020<br>Horstmann et al. 1993<br>Sharan et al. 2021 |
| **13** | **20.9** | **~24**<br>**~20-25**<br>**~21** | **β-legumin** | Warsame et al. 2018<br>Tucci et al. 1991<br>Ashraf et al. 2020<br>Horstmann et al. 1993<br>Sharan et al. 2021 |
| **14** | **< 20.0** | **~20** | **β-legumin** | Tucci et al. 1991<br>Horstmann et al. 1993<br>Sharan et al. 2021 |
| 15 | < 20.0 | - | - | - |
| 16 | < 20.0 | - | - | - |
| 17 | < 20.0 | - | - | - |

*Note:* Protein band numbers 14 to 17 are given as < 20.0 kDa. This is, since some gels contained as smallest marker protein the 20 kDa band and the imaging software would not extrapolate below 20 kDa.

Via metabolic fingerprinting, 5598 metabolic features were found, with the number of features per sample ranging from 1455 to 2285 (Figure 3c). Of these metabolic features, 202 (i.e., 3.6%) were found in all samples. There was a tendency for a positive relationship between the number of metabolic features and the protein content, but it was not significant (Spearman correlation: *ρ* = 0.15, *p* = 0.077). In contrast, the number of metabolic features showed a significant negative correlation with the sulfur content (Spearman correlation: *ρ* = -0.19, *p* < 0.05) of the seeds. The number of metabolic features mainly differed between field sites, with Scharnhorst and Nossen showing the highest numbers, while Astrup and Rauischholzhausen had the lowest numbers. The distribution of metabolite patterns over the samples (NMDS, stress value: 0.176) was similar to those observed for the proteins (Figures 4a,b; Mantel test: *r* = 0.33, *p* < 0.001). As for the protein patterns, the metabolite patterns were strongly cultivar-specific. The cultivars C06 and C10 were most distinct from each other, with C10 showing one outlier (Figure 4b, left). Several other cultivars were separated from each other as well. Compared to the protein patterns, the metabolite patterns showed more variation between the field sites (Figure 4b, right). The most obvious differences were found between Scharnhorst and Astrup. The NMDS ordination of metabolite patterns was not correlated with the sulfur content, but with the protein content as well as with the number of metabolic features (Figure 4b, second line). The protein content varied along the separation of cultivars, from higher values in the cultivar C06 to lower values in C03, but showed some variation along with the separation of field sites as well. In contrast, the number of metabolic features correlated mainly with the separation of the field sites, from higher numbers of features in samples from Scharnhorst to lower numbers in those from Astrup.

## 4 Discussion

This study revealed that the seed chemistry of faba bean is highly diverse and variable, with the protein content, sulfur content, several proteins as well as manifold metabolites differing between cultivars and/ or field sites.

The seed protein contents were generally very high, with up to 33.6% dry mass. This is in line with other studies showing that seed protein contents of *V. faba* exceed those of cereal grains by far, but that they may also be higher than in other grain legumes (Rahate et al. 2021; Sharan et al. 2021; Walter et al. 2022). The seed sulfur contents were in the range reported previously for various faba bean cultivars (Bhatty, Finlayson, and Mackenzie 1977; Makkar et al. 1997). A large part of the sulfur was probably due to the sulfur-containing amino acids methionine and cysteine (Bhatty et al. 1977).

We assume that specialized metabolites did not contribute much to the sulfur content, because most specialized metabolites, at least in plant taxa other than Brassicaceae and Alliaceae, do not contain sulfur (Abdalla and Mühling 2019; Hill et al. 2023). The positive correlation between the protein content and sulfur content we found does not correspond to a study showing no correlation between these traits for six *V. faba* cultivars (Bhatty et al. 1977), but is in line with a study including 35 faba bean genotypes (Warsame et al. 2020).

Based on comparison with published SDS-PAGE patterns, the major protein bands found in the *V. faba* seeds were assigned as globulins of the vicilin/ convicilin type as well as of the legumin type.

Indeed, the globulins are the major storage proteins in faba bean seeds, where these salt-soluble proteins are comprising approximately 85% of the total protein (Sharan et al. 2021). We did not assign the bands of minor intensity in our SDS-PAGE analysis to specific bands in published patterns. However, workers that examined total lanes of SDS-PAGE-separated proteins from *V. faba* seeds with mass spectrometric techniques reported identification of further vicilin/ convicilin and legumin type subunits as well as metabolically active proteins (Liu et al. 2017; Warsame et al. 2020). Thus, it has to be expected that our SDS-PAGE results contain a similar protein pattern, particularly since we followed the extraction procedure of Warsame et al. (2020). It should also be noted that most bands will contain more than one protein, as has been specifically found by Warsame et al. (2020).

In addition, we detected various metabolic features, probably comprising the manifold metabolites from different classes described for *V. faba* seeds (Abu-Reidah et al. 2014; Mekky et al. 2020; Valente et al. 2019). The correlation of the protein patterns with the metabolite patterns indicates that proteins are tightly linked with metabolism. While most proteins in mature faba bean seeds are storage proteins and enzymatically inactive (Sharan et al. 2021; Warsame et al. 2022), some enzymes may have shaped the metabolome, probably mainly in earlier stages of seed development. Moreover, the competition of proteins and metabolites for (limited) resources probably explains the tight link between the seed protein and metabolite patterns.

The high cultivar specificity of the seed protein content, protein and metabolite patterns in our study is remarkable. As *V. faba* is partially allogamous, with out-crossing rates depending on the genotypes and environment (Gnanasambandam et al. 2012), we cannot rule out that some cross-pollination between different cultivars occurred within the field sites, exceeding natural levels due to the small sizes of the subplots compared to large agricultural fields. However, we assume that most seeds were not derived from between-cultivar crossings and thus represent the assigned cultivar. In line with our study, many other studies reported cultivar specificity in seed protein contents (e.g., Khazaei and Vandenberg 2020; Makkar et al. 1997; Mayer Labba et al. 2021; Mohammadi et al. 2024; Segers et al. 2022). Such differences are probably largely due to breeding, either directly for high protein contents or indirectly for other traits that also affect protein contents. For example, a negative relationship between yield and protein content has been reported for 17 *V. faba* cultivars at Nordic locations (Skovbjerg et al. 2020). Moreover, negative relationships between seed tannin and protein contents occur (Khazaei and Vandenberg 2020; Walter et al. 2023). The composition of proteins and metabolites in *V. faba* seeds has been less investigated in comparative studies including different faba bean cultivars until now. However, several cultivars were included in investigations of the seed protein (Alghamdi 2009; Warsame et al. 2020) and metabolite (Mekky et al. 2020; Valente et al. 2019) composition, mainly focusing on the characterization of (new) proteins and metabolites.

There were indications that 14 *V. faba* cultivars grown in Sweden varied in several seed nutrients and antinutrients (Mayer Labba et al. 2021). The metabolic composition of faba bean seeds was also interactively affected by seed traits (color, weight) and genotypes (cultivated versus landraces; Choi et al. 2024). Moreover, in a comparative metabolomics approach with seeds of two *V. faba* genotypes, circa 150 differently abundant metabolites were found, with flavonoid biosynthetic pathways being most distinct (Shi et al. 2022). As discussed for the protein content, the cultivar specificity of protein and metabolite patterns in our study is probably a direct or indirect result of breeding. Remarkably, our study indicates that cultivar differences in seed chemistry go far beyond breeding targets like the protein content or certain metabolites (vicine/ convicine, tannins). Next to protein-metabolite links and limited resources, complex connections within metabolic networks probably contribute to these far-reaching effects. For example, tannins may bind to proteins and modify their activity and function (Crépon et al. 2010; Wink 2003), which may in turn affect other metabolites, contributing to metabolic differences between low- and high-tannin cultivars going beyond the tannin pathway.

Several chemical traits of the faba bean seeds showed variation between the field sites. While the protein content and patterns were less affected, the sulfur content, number of metabolic features and metabolite patterns largely varied between field sites. The differences between field sites did not follow an obvious geographic pattern (Figure 1), indicating that probably not only climatic factors with geographic gradients (e.g., temperature, precipitation) but also other environmental variables contributed. Likely, several (a)biotic factors acted in concert. For example, soil types and farming practices affecting the availability of nutrients and water may have impacted protein biosynthesis and metabolic pathways relying on these resources. Another study showed that, under real farming conditions with several *V. faba* cultivars all over Germany, the seed protein contents were affected by the cultivation year, partly correlating with temperature, precipitation and soil composition (Walter et al. 2022). Moreover, a tendency for higher seed protein contents at lower irrigation was found in a study with 13 faba bean genotypes (Alghamdi 2009). In a controlled experiment, seed maturation was delayed at lower temperatures, with implications for contents of starch and carbohydrates (Lundby et al. 2023). The large differences in seed sulfur content between field sites may reflect variation in the availabilities of sulfur-containing nutrients, being related to soil type, cultivation history and/ or fertilization. Within the crop plant taxa, Fabaceae require less sulfur than Brassicaceae, but more than Poaceae (Hawkesford et al. 2023). Sulfur fertilization led to higher contents of sulfur, proteins and tannins in faba bean seeds, similar to nitrogen fertilization (Elsheikh and Elzidany 1997).

Biotic interactions may have further shaped the seed chemistry. For example, *V. faba* cultivars differ in traits related to rhizobia-mediated N_2_-fixation (Abu et al. 2024; Kilian et al. 2001), while the availability of symbiotically effective rhizobia is decisive and may depend on the field site (Allito, Ewusi-Mensah, and Logah 2020; Denton, Pearce, and Peoples 2013). N_2_-fixation is also influenced by the availability of certain elements including sulfur (Becana, Wienkoop, and Matamoros 2018; Hawkesford et al. 2023). In turn, N_2_-fixation has probably large influences on proteins and metabolites, as nitrogen is part of many proteins including enzymes and of metabolites. Similarly, plant antagonists may have contributed to cultivar and field site differences. Indeed, faba bean cultivars vary in their susceptibility to antagonists (e.g., *Bruchus* spp. and leaf pathogens: Khazaei et al. 2021; Maalouf et al. 2019; Segers et al. 2022) and field sites may differ in the occurrence of these antagonists. Although we only analyzed seeds that were not infested with *Bruchus* spp. and many of the pathogens are only found at the leaves, such antagonists may have affected the proteins and metabolites systemically in (developing) seeds.

In our study, most chemical traits of *V. faba* seeds showed a higher consistency between field sites than between cultivars. This indicates that the investigated cultivars are distinct and more or less stable in different environments. Similar to (yield) stability, which is often considered in breeding and selection of cultivars (Maalouf et al. 2019; Mohammadi et al. 2024), stability in seed quality is of major importance for the market. We included only commercial spring cultivars available on the German market, which were grown on field sites with conventional agriculture in one growing season, and we discarded unusual, infected and damaged seeds. Inclusion of more cultivars and field sites, especially in other countries with distinct environmental conditions, and more growing seasons would probably lead to an even broader spectrum of seed chemistry. Likewise, seed chemistry may vary between spring and winter cultivars and between fields with conventional versus organic agriculture, as demonstrated for protein contents of grain legumes (Segers et al. 2022; Walter et al. 2022). While seeds with a high infestation (e.g., with *Bruchus* spp.) may not be placed on the market, a certain, low degree of infestation may be accepted, but probably further affects the seed quality.

From a nutritional and health viewpoint, the results of our study are of high importance, as both proteins and metabolites contribute to the quality of faba bean seeds (Feng et al. 2024; Martineau- Côté et al. 2022). While the high protein contents underline the value of *V. faba* for nutrition, the protein composition has to be considered as well. Compared to cereal grains, faba bean proteins are rich in lysine, but poor in tryptophan and sulfur-containing amino acids (Crépon et al. 2010; Sharan et al. 2021). Sulfur compounds are important for the biosynthesis of several compounds in humans, with methionine being essential and cysteine semi-essential (Hill et al. 2023). Under sulfur deficiency, sulfur-poor proteins may be up-regulated at the expense of sulfur-rich ones (Aarabi et al. 2020). For faba bean, where globulins contain less sulfur than other proteins (Sharan et al. 2021), a shift to more globulins would exacerbate the limitation regarding methionine and cysteine despite high protein contents. We found a huge variation in seed sulfur contents related to field sites. However, as we did not measure sulfur-containing amino acids in the proteins, it is unclear whether this variation affected the nutritional quality. Based on the critical role of the sulfur-containing amino acids in seeds, efficient sulfur fertilization should be assured in faba bean farming. Our study also revealed strong metabolic differences between cultivars and field sites, going far beyond single metabolites targeted during breeding. The identification of metabolites was beyond the scope of our study, but future studies may shed more light on these metabolites. To evaluate the relevance of varying protein and metabolite compositions for nutrition and health, the interplay between compounds should also be investigated, for example with regard to protein digestibility and the bioavailability of minerals.

Furthermore, as processing (e.g., soaking, dehulling, cooking) of faba bean seeds influences their chemical composition (Badjona et al. 2023; Corzo-Ríos et al. 2022; Sharma and Sehgal 1992), processing steps should be considered in the evaluation of the seed quality of this species.

## 5 Conclusions

Taken together, our study highlights that breeding of faba bean for certain traits and cultivation at different field sites had far-reaching impacts on seed chemistry, both at the protein and the metabolite level. More comprehensive studies are needed to better understand the variation in seed quality of this important sustainable crop species and the implications for the nutrition and health of livestock and humans. Finally, research may assist breeders with breeding and selection of cultivars and farmers with choosing appropriate field sites to use the full potential of faba bean.

## Supporting information

Graphical Table of Contents

## Author Contributions

**Julia Brandt:** investigation, formal analysis, writing – review & editing, visualization. **Alexandra Hüsken:** resources, conceptualization, methodology, validation, investigation, writing – review & editing. **Eileen Whitelaw:** methodology, investigation, formal analysis, writing – review & editing. **Georg Langenkämper:** conceptualization, methodology, validation, investigation, formal analysis, writing – review & editing, visualization, supervision, project administration. **Rabea Schweiger:** conceptualization, methodology, validation, investigation, formal analysis, writing – original draft, visualization, supervision, project administration.

## Acknowledgments

The metabolomics analyses were supported by basic funding of the Department of Chemical Ecology provided by Bielefeld University/ Faculty of Biology. Determination of protein and sulfur content as well as protein analyses were supported by basic funding of the Department of Safety and Quality of Cereals, MRI. We thank the Bundessortenamt for providing seeds, Caroline Müller for valuable input and discussion as well as Sophie Beyer, Lidia Arent, Fiona Ott, Gabriele Ortmann, Annette Meyer-Wieneke and Sascha Schulz for practical help.

## Conflicts of Interest

The authors declare no conflicts of interest.

## Data Availability Statement

Data will be made available upon reasonable request.

## Funding

The metabolomics analyses were supported by basic funding of the Department of Chemical Ecology provided by Bielefeld University/ Faculty of Biology. Determination of protein and sulfur content as well as protein analyses were supported by basic funding of the Department of Safety and Quality of Cereals, MRI.

## Conflicts of Interest

The authors declare no conflicts of interest.

## Ethics Approval Statement

Not applicable.

## Patient Consent Statement

Not applicable.

## Permission to Reproduce Material from Other Sources

Not applicable.

## Notes

### Competing Interest Statement

The authors have declared no competing interest.

