## Supplementary material for "Cultivar- and field site-specific protein and metabolite patterns of faba bean (*Vicia faba* L.) seeds": Graphical Table of Contents

In our study on the seed chemistry of faba bean (*Vicia faba* L.), the protein content as well as the protein and metabolite patterns largely differed between 11 cultivars and were less affected by the field site. In contrast, the seed sulfur content depended mainly on the field site. Our study shows that breeding has led to far-reaching chemical changes in faba bean seeds, which is important in the light of their nutritional quality for livestock and humans.

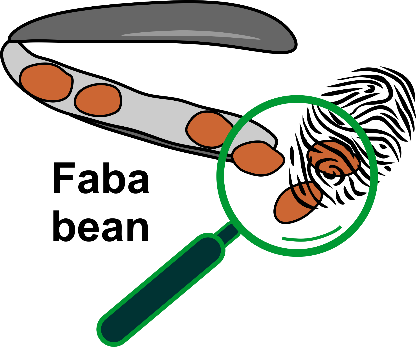
